# Deficiency of complement component C1Q prevents cerebrovascular damage and white matter loss in a mouse model of chronic obesity

**DOI:** 10.1101/2020.02.03.931949

**Authors:** Leah C. Graham, Heidi E. Kocalis, Ileana Soto, Gareth R. Howell

## Abstract

Age-related cognitive decline and many dementias involve complex interactions of both genetic and environmental risk factors. Recent evidence has demonstrated a strong association of obesity with the development of dementia. Furthermore, white matter damage is found in obese subjects and mouse models of obesity. Here, we found that components of the complement cascade, including C1QA and C3 are increased in the brain of western diet (WD)-fed obese mice, particularly in white matter regions. To functionally test the role of the complement cascade in obesity induced brain pathology, female and male mice deficient in complement component 1qa (C1QA), an essential molecule in the activation of the classical pathway of the complement cascade, were fed a WD and compared to WD-fed WT mice, and to *C1qa* knockout (KO) and WT mice fed a control diet (CD). *C1qa* KO mice fed a WD became obese but did not show pericyte loss or a decrease in laminin density in the cortex and hippocampus that was observed in obese WT controls. Furthermore, obesity-induced microglia phagocytosis and breakdown of myelin in the corpus callosum were also prevented by deficiency of C1QA. Collectively, these data show that C1QA is necessary for damage to the cerebrovasculature and white matter damage in diet-induced obesity.

**SIGNIFICANCE STATEMENT:** Economic growth, an increasingly sedentary lifestyle and a nutritional transition to processed foods and high calorie diets have led to a significant increase in obesity prevalence. Several chronic diseases have been associated with obesity, including dementia. Obesity-induced, peripheral inflammation has been proposed as a possible trigger of pathological changes in the brain that lead to cognitive dysfunction and predisposition to dementia. Here we show that genetic deletion of the complement component C1QA prevents cerebrovascular damage, neuroinflammation and white matter degradation in a mouse model of western diet-induced obesity, demonstrating that inflammatory responses play a significant role in obesity-induced brain pathology. The complement pathway is an attractive therapeutic target to prevent cognitive decline and reduction of dementia risk caused by obesity.

## INTRODUCTION

Over the last 30 years obesity prevalence has significantly increased in many countries to the point that one third of the world population is overweight or obese (Hruby and Hu, 2015). In fact, it is projected that in the USA over 85% of adults will be overweight or obese by 2030 (Wang et al., 2008; Hruby and Hu, 2015). Obesity has been defined as an excess of body weight for height (body mass index >29). However, because of its metabolic alterations and increased risk for the development of chronic diseases, obesity is now considered an energy metabolism disorder (Hruby and Hu, 2015). Moreover, obesity has also been associated with the development of mental disorders, cognitive dysfunction and dementia (Elias et al., 2005; Viscogliosi et al., 2013; Singh-Manoux et al., 2018). Little is known about how obesity promotes pathological changes in the brain that are not necessarily associated to chronic conditions such as metabolic syndrome and Type 2 diabetes. Nonetheless, obesity-induced inflammation has been identified as a possible trigger of pathological changes in the brain that lead to cognitive dysfunction and predisposition to neurodegenerative diseases like multiple sclerosis and Alzheimer’s disease (AD) (Miller and Spencer, 2014).

Previous studies in our laboratory have shown that western diet (WD)-induced obesity amplifies neuroinflammation in the cortex and hippocampus of aged mice and in a mouse model of AD (Graham et al., 2016; Graham et al., 2019). More recently, we found significant infiltration of peripheral myeloid cells and neutrophils in the brain of diet-induced obese mice, particularly in the cortex, the hippocampus and the corpus callosum (Yang, 2019). These strong neuroinflammatory responses are accompanied by behavioral deficits, neurovascular decline and increased degradation of myelin by resident microglia or peripherally-derived myeloid cells, despite the increased expression of myelin proteins by oligodendrocytes (Graham et al., 2016; Graham et al., 2019). Importantly, white matter integrity throughout the corpus callosum and frontal white matter is reduced in cognitively normal older adults, patients with mild cognitive impairment (MCI) and AD patients who are obese (Graham et al., 2019), suggesting that white matter damage is a pathological hallmark of obesity.

It can be presumed that the chronic low-grade inflammatory state in obesity influences the neurovascular decline and the activation of microglia observed in the brain leading to subsequent phagocytosis of myelin, but this has not been critically tested. Furthermore, *in vitro* studies have shown that myelin opsonization with complement components and the presence of the complement receptor CR3 by microglia are required for maximal phagocytosis of myelin, suggesting an important role of the complement pathway in myelin phagocytosis (DeJong and Smith, 1997). It has been shown that in multiple sclerosis lesions complement components colocalize with areas of active myelin degradation along with the increased density of microglia/macrophages expressing complement receptors (Barnett et al., 2009; Grajchen et al., 2018; Loveless et al., 2018). Although, it is known that several components of the classical complement pathway are produced by adipose and peripheral immune cells in obese mice (Zhang et al., 2007), evidence of activation of the complement pathway in the brain during obesity is lacking. Here, we hypothesized that myelin phagocytosis by microglia (or peripherally-derived myeloid cells) in WD-induced obese mice was mediated by the activation of the classical complement pathway. We found that genetic deletion of *C1qa* did not change the body composition or common blood-based markers of metabolic syndrome in the western diet-induced obese mice. However, C1QA deficiency did significantly lessen cerebrovascular damage and the activation and phagocytic activity of microglia in the cortex, hippocampus and corpus callosum preventing the degradation of myelin.

## MATERIALS AND METHODS

### Animals

All methods are in accordance with The Jackson Laboratory Institutional Animal Care and Use Committee (IACUC) approved protocols. C57BL/6J (B6) mice (JAX stock # 000664), B6.C1qa^tm1a(EUCOMM)Wtsi^/J, and B6.C3^tm1Crr^/J (Jax stock #003641) mice were used in this study and maintained in the Howell Lab colony. B6.C1qa^tm1a(EUCOMM)Wtsi^/J mice were created by backcrossing B6N.C1qa^tm1a(EUCOMM)Wtsi^/J mice at least 10 generations to C57BL/6J. Males were initially used exclusively in this study to avoid effects of the estrus cycle, but a second cohort of male and female mice were used to determine possible sex differences in response to both diet and complement deficiency. All mice were maintained on a 12/12 hours (hrs) light/dark cycle. Cohorts were maintained from wean on standard LabDiet® 5K52 (referred to as “Control diet” or “CD”). Half of the cohorts were switched to TestDiet® 5W80 (referred to as “Western diet” or “WD”) (Graham et al., 2016).

### Metabolic profiling

#### Glucose and insulin tolerance test

Glucose and insulin tolerance tests were performed in 12 mos mice, following 10 mos on the WD. After fasting for 5 hours, glucose tolerance test (GTT) was performed (Agri Laboratories, LTD). For the insulin tolerance test (ITT), mice received a single intraperitoneal injection of a diluted insulin solution (males 0.75 IU/kg and females 0.50 IU/kg; prepared in sterile saline) (Humulin R U-100, Eli Lilly). Glucose was determined using a handheld glucometer (Bayer Contour Next).

#### NMR Body Composition

Body composition was determined in the Echo MRI 3-in-1, time domain nuclear magnetic resonance (TD-NMR) system (Houston, TX). Body weight was measured on a standard laboratory balance before mice were placed in to a clear, cylindrical holder. The tube was gently inserted in to the boor for an approximately two-minute measurement. Adiposity (% body fat) was calculated as ((fat mass/total body weight) x 100).

#### Plasma lipid measurements

Blood was collected in K2 EDTA (1.0mg) microtainer tubes (BD) at harvest (non-fasted) and kept at room temperature for at least 30 minutes to prevent clotting and then centrifuged at 22°C for 15 minutes at 5000rpm. Plasma was carefully collected and aliquoted. Plasma was characterized on the Beckman Coulter AU680 chemistry analyzer.

### Behavioral battery

All the tasks performed by The Jackson Laboratory’s Mouse NeuroBehavioral Facility (MNBF) were previously validated using control B6 mice. The *spontaneous alternation* task, a measure of working memory, was conducted as previously described (Sukoff Rizzo et al., 2018). Briefly, a clear Plexiglas Y-maze was used under adjusted, ambient lighting (∼ 50 lux). Subject mice were acclimated to the testing room for 1hr prior to testing. Tested mice were then placed midway of the start arm (A), facing the center of the Y for an 8-minute test period, and the sequence of entries into each arm are recorded via a ceiling mounted camera integrated with behavioral tracking software (Noldus Ethovision). The percentage of spontaneous alternation was calculated as the number of triads (entries into each of the 3 different arms of the maze in a sequence of 3 without returning to a previously visited arm) relative to the number of alteration opportunities.

For assessment of *open field activity*, Open Field Arenas (40 cm × 40 cm × 40 cm; Omnitech Electronics, Columbus, OH) were used. A light fixture mounted ∼50 cm above the center of each arena provided a consistent illumination of ∼400–500 lux in the center of the field. Before the test, mice were acclimated to an anteroom outside the testing room for a minimum of 1 hour. Subsequently, the tested mice were placed individually into the center of the arena where the infrared beams recorded distance traveled (cm), vertical activity, and perimeter/center time.

For *grip strength*, subjects were weighed and acclimated for at least 1hr prior to the test. All the grip strength tests were assessed using the Bioseb grip strength meter equipped with a grid suited for mice. For forepaw and four paw grip strength testing, mice were lowered towards the grid by their tails to allow for visual placing and for the mouse to grip the grid with their paws. Subjects were firmly pulled horizontally away from the grid (parallel to the floor) for 6 consecutive trials with a brief (<30 sec) rest period on the bench between trials. Trials 1-3 tested only the forepaw grip; while the following trials 4-6 included all four paws. The average of the 3 forepaw trials and the average of the 3 four paw trials were analyzed with and without normalization for body weight.

### Mouse perfusion and tissue preparation

Tissue was collected at 3.5 mos, 12 mos, and 20 mos (aged C3 mice). Mice were anesthetized with a lethal dose of ketamine/xylazine, transcardially perfused with 1X phosphate buffered saline (PBS) and brains carefully dissected and hemisected in the midsagittal plane. One half was snap-frozen, the other half immersion fixed in 4% paraformaldehyde (PFA) for two nights at 4°C. After fixation, half brains were rinsed in 1 x PBS, immersed in 30% sucrose (in 1 x PBS) overnight at 4°C before being frozen in Optimal Cutting Temperature (OCT) compound. Tissue was cryosectioned at 25μm on to glass slides for RNA in situ hybridization and immunofluorescence.

### RNA in situ hybridization

For in situ hybridization, an RNA probe for *C1qa* (GE Dharmacon Clone ID 3592169) was synthesized, labeled with digoxigenin (Dig) and hydrolyzed by standard procedures. Frozen sections were post fixed (4%PFA for 5min), rinsed twice with 1X PBS and acetylated with 0.25% acetic anhydride for 10min in 0.1M triethanolamine (TEA). Sections were then washed in PBS and incubated overnight at 65°C in hybridization solution [50% formamide, 1X Hybe solution (Sigma-Aldrich), 1mg/ml yeast RNA] containing 1g/ml Dig-labeled riboprobe. After hybridization, sections were washed by immersion in 0.2X saline-Sodium citrate buffer at 72 °C for 1hr. Dig-labeled probes were detected with an AP-conjugated anti-Dig antibody (Roche) followed by NBT/BCIP (nitroblue tetrazolium/5-bromo-4-chloro-3-indolyl phosphate) reaction (Roche). After *in situ* hybridization, sections were incubated with DAPI for nuclei staining and mounted in Aqua PolyMount (Polysciences) as described previously (Howell et al., 2011).

### Immunofluorescence

For immunostaining with antibodies against vascular associated proteins, sections were pretreated with pepsin as previously described (Soto et al., 2015) with minor modifications. After pepsin pre-treatment, sections were incubated overnight with the following primary antibodies: goat anti-PDGFRβ (1:40, R&D), and rabbit anti-LAMININ (LAM,1:200, Sigma-Aldrich). Sections used for non-vascular associated protein visualization were incubated overnight in primary antibodies: rabbit anti-MBP (1:200, Abcam), or rat anti-MBP (1:200, Abcam), goat anti-IBA1 (1:100, Abcam), or rabbit anti-IBA1 (1:100, Wako), rat anti-CD68 (1:100, Bio-Rad), rabbit anti-C1QA (1:100, Abcam), and goat anti-C3 (1:75, R&D systems). After primary antibodies incubation, the sections were washed 1X PBT (1X PBS + 1% Triton100X) and incubated with secondary antibodies for 2 hours at RT. After incubation with secondary antibodies, sections were stained with DAPI and mounted in Aqua PolyMount (Polysciences).

### Imaging and quantification

#### Imaging

For each mouse, four images per brain region (parietal cortex, corpus callosum and CA1 region of the hippocampus) were generated. Quantifications were performed per image – therefore where images contained multiple brain regions (e.g. parietal cortex, corpus callosum and CA1 region of the hippocampus) these were batched together (see *Experimental design and statistical analysis* for more details). For quantifying cell number or area, images were captured on a Zeiss AxioImager microscope. For each antibody, all images were captured using identical parameters for accurate quantification. Where possible, fluorescent intensity was standardized to samples from CD-fed mice. However, given the striking difference in intensity between CD-fed and WD-fed mice for MPB, images were standardized to WD-fed mice.

#### Quantification in FIJI (ImageJ)

Images for IBA1^+^DAPI^+^ cells were manually counted using the cell counter plugin FIJI v1.0. For quantification of PDGFRβ, and LAMININ, fluorescent area was calculated using a previously validated in-house Vascular Network Toolkit (VNT) plugin for FIJI v1.0 (Soto et al., 2015). This segmentation algorithm was written as an ImageJ macro for automated processing of images that includes the following steps: Gaussian blur (2 px), “Find edges”, Variance (5 px), Median (3 px), Subtract (σ px), Multiply (255), Invert and Analyze Particles. The analysis of all the images was automated and blind to the experimenters.

#### Quantification and visualization in IMARIS

Quantitative analysis of microglia activation (IBA1^+^ cells), microglia phagocytic activity (CD68^+^ phagosomes), and myelin degradation (MBP) were performed using IMARIS 8.1 (Bitplane) as previously described (Graham et al., 2019). Briefly, images captured on the Leica SP5 confocal microscope. Z stacks were compiled with 0.2 μm steps in the z direction with 1024×1024 pixel resolution. For each antibody, all images were transferred to IMARIS and rendered using identical parameters and the co-localization tool was used to determine both the surface areas of MBP, IBA1 and CD68 and the interactions between surfaces.

### Experimental design and statistical analysis

For behavioral analyses, groups of 10-12 mice per sex per genotype per diet allowed us to detect effects of greater than 1.15 standard errors at 80% power as shown previously (Graham et al., 2019). Changes to vascular structures, white matter and microglia were identified using sample sizes of at least 5. Samples sizes are provided in the figure legends (n=biological replicate and refers to number of mice/samples used in each experiment). Region of interest was standardized for all statistical analyses: Hippocampus CA1 (region including stratum radiatum, pyramidal layer and stratum oriens), Cortex (region including posterior cortex, layers 4-6), Corpus Callosum (region between CA1 and posterior parietal cortex). Regions determined based on Allen Brain Atlas.

All data were analyzed using GraphPad Prism software. For comparisons between two groups, significance was calculated using unpaired t-tests. For multiple comparisons, significance was calculated using one-way multifactorial analysis of variance (ANOVA) followed by Tukey posthoc tests. ANOVA assumes equal variance and this was determined prior to each test using the Brown-Forsythe test. *p*-values provided are as stated by GraphPad Prism software and significance was determined with *p*-values less than 0.05. Standard error of the mean was used in all graphs.

## RESULTS

Transcriptional profiling demonstrated that WD-induced obese male mice have an increased expression of complement components, including the central components *C1qa*, *C1qb*, *C1qc* and *C3*, in the brain when compared to non-obese mice fed a control diet (CD) (**Fig. 1A**, (Graham et al., 2019)). To determine the location and extent of complement deposition in the brain of obese mice, male mice were fed either a WD or CD from 2 months of age (mos) for 10 months. RNA in situ hybridization in 12 mos brains showed a significant increase in the numbers of *C1qa*+ expressing cells in the cortex and hippocampus from WD-fed compared to CD-fed mice (**Fig. 1B-C**). Also, the intensity and size of staining for each *C1qa+* cell appeared greater in WD-fed compared to CD-fed mice, features previously suggested to represent activated myeloid cells (Davis et al., 2017). C1Q and C3 immunostaining appeared greater in the corpus callosum in WD-fed compared to CD-fed mice (**Fig. 2**). Little to no C1Q or C3 immunostaining was observed in the corpus callosum of young mice or in brains from mice deficient in *C1qa* and *C3* (**Fig. S1**). The increase in complement components in WD-fed compared to CD-fed mice appeared due to changes in microglia as C1Q and C3 immunoreactivity co-localized with IBA1 – a marker of microglia.

**Figure 1.**
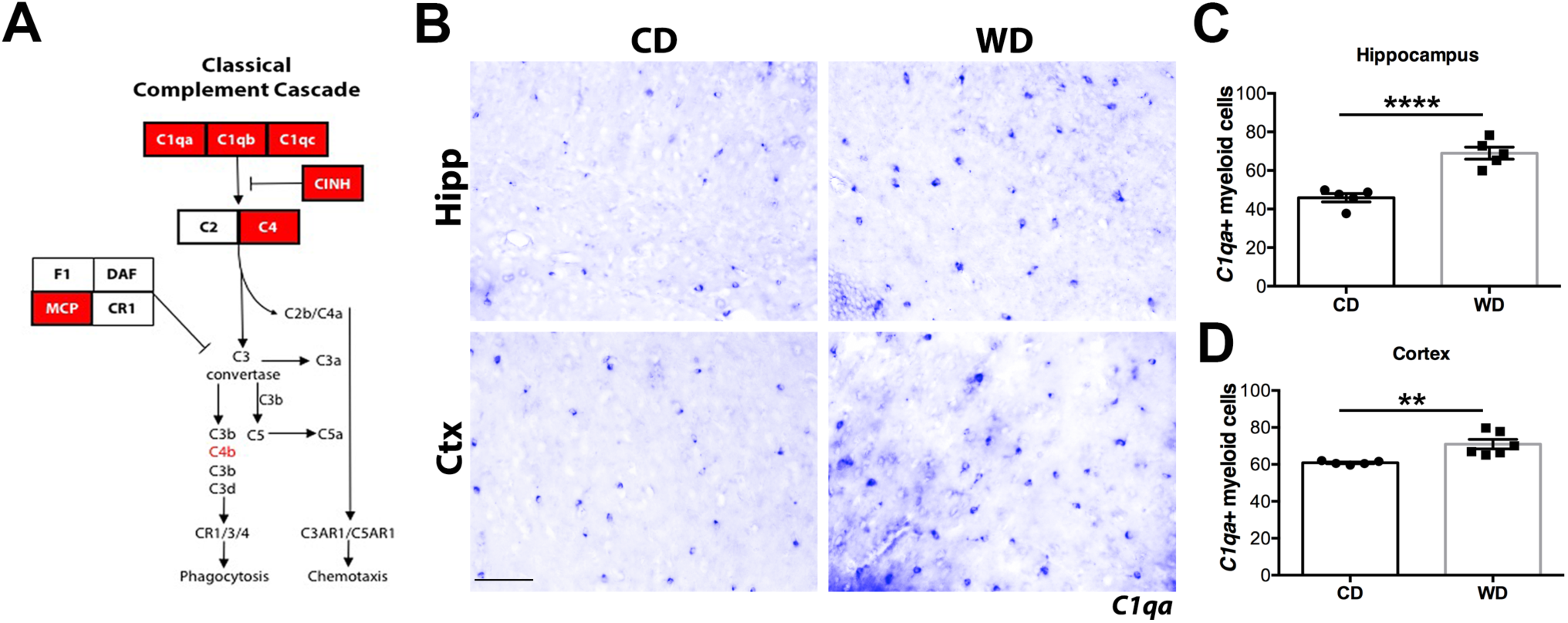
Complement components are increased in brains of WD-fed obese mice. (**A**) Expression of complement proteins upregulated in brains of WD-fed mice (red boxes). (**B-D**) Increased number of *C1qa+* cells in the hippocampus and cortex of WD-fed mice. Data are presented as mean ± SEM, n ≥ 5, **P < 0.01 and ****P < 0.001 by paired t-test. Scale bars: B, 50μm.

**Figure 2.**
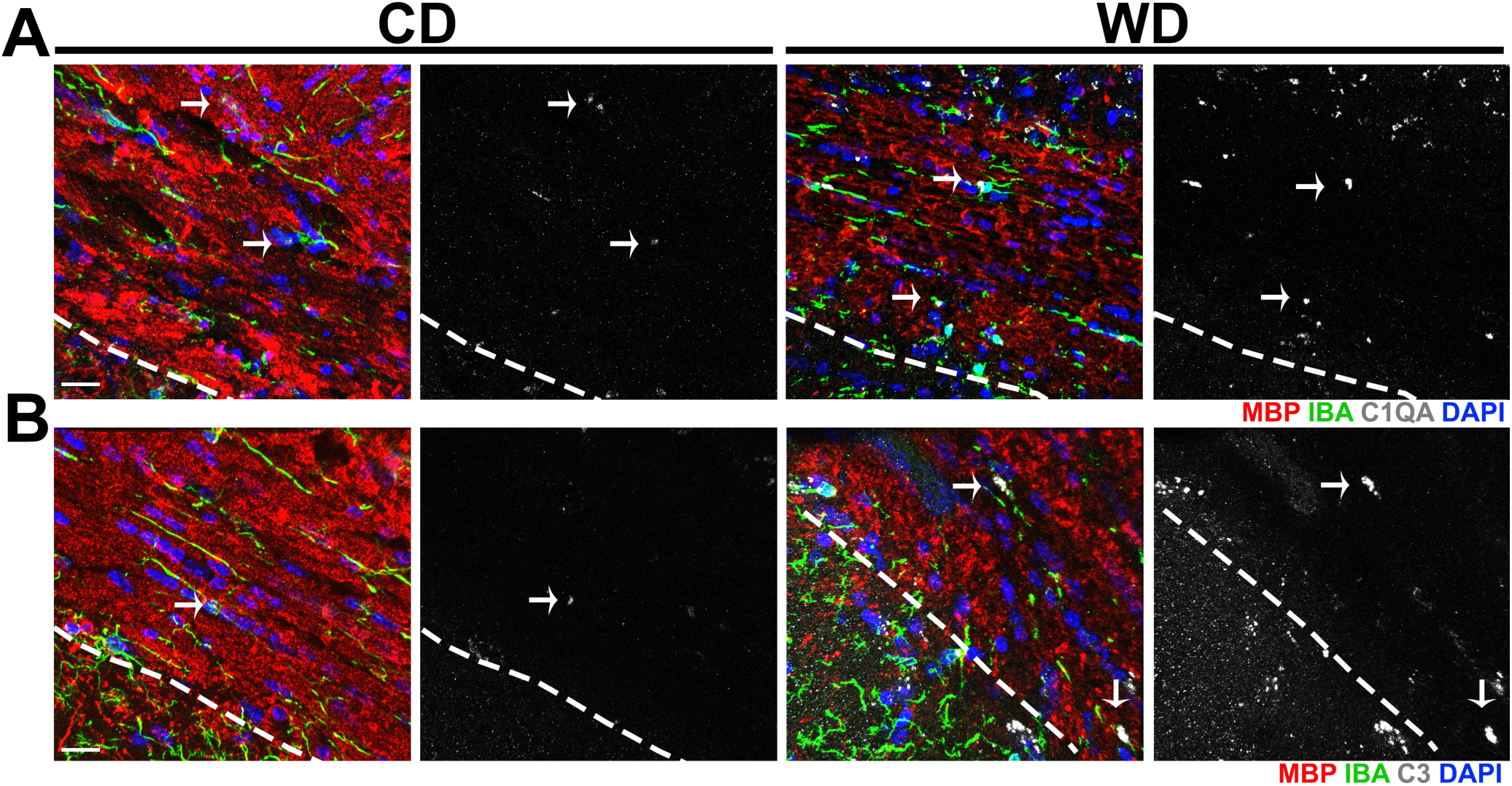
C1QA and C3 deposits increase in white matter tracts of WD-fed mice. (**A**) Increased C1QA immunoreactivity in the corpus callosum of WD-fed compared to CD-fed mice colocalized with IBA1^+^ cells and MBP^+^ myelin. (**B**) Increased C3 immunoreactivity in the corpus callosum of WD-fed compared to CD-fed mice colocalized with MBP^+^ myelin. White dotted lines indicate the boundary between the CA1 region of the hippocampus and the corpus callosum. Scale bars: A-B, 40μm.

To elucidate the role of complement activation in brains of obese mice, B6.*C1qa^-/-^*(*C1qa* KO) mice were generated (see methods, **Fig. S2A**). RNA in situ hybridization with *C1qa* riboprobes confirmed the complete absence of *C1qa* expression in the brain of B6.*C1qa^-/-^* mice (**Fig. S2B, C**). Cohorts of 10-12 male and female *C1qa* KO and B6 (WT) control mice were established. At 2 mos, mice were fed either a WD or a CD. At 12 mos, mice were assessed for characteristics of metabolic syndrome that included weight, glucose and insulin tolerance. WD-fed *C1qa* KO male and female mice both gained weight similarly to their WD-fed WT littermate controls (**Fig. 3A, C**). Consistent with higher total weight, the percentage of adiposity was greater in males and females from both WD-fed mouse strains when compared to CD-fed mice (**Fig. 3B, D**). Lipid profiling of plasma from WD-fed and CD-fed mice showed that total cholesterol levels in males and females were increased in both *C1qa* KO and WT mice when compared to CD-fed mice (**Fig. 3E, G**). Triglyceride levels were decreased in *C1qa* KO males and unchanged in females from WT and C1QA deficient mice (**Fig. 3F, H**). The levels of LDL were only increased in males by the WD in both mouse strains (**Fig. 3I, K**), while HDL levels were increased in both females and males from WD-fed mice (**Fig. 3J, L**). Changes in the level of fatty acids were not found in any of the experimental groups (**Fig. 3M, N**). Obesity is a high risk for insulin resistance, therefore glucodynamic responses were evaluated in WD-fed and CD-fed WT and C1QA deficient mice. While no changes in blood glucose levels were found in the basal, non-fasting state (**Fig. 4A, C**), only WD-fed WT males presented a moderate impaired glucose tolerance, that was prevented by the deficiency of C1QA in the WD-fed *C1qa* KO males (**Fig. 4B, D**). Finally, no differences in insulin levels were observed (**Fig. 4E-H**). Collectively these data show WD-fed *C1qa* KO and WT mice were obese but not insulin resistant or diabetic.

**Figure 3.**
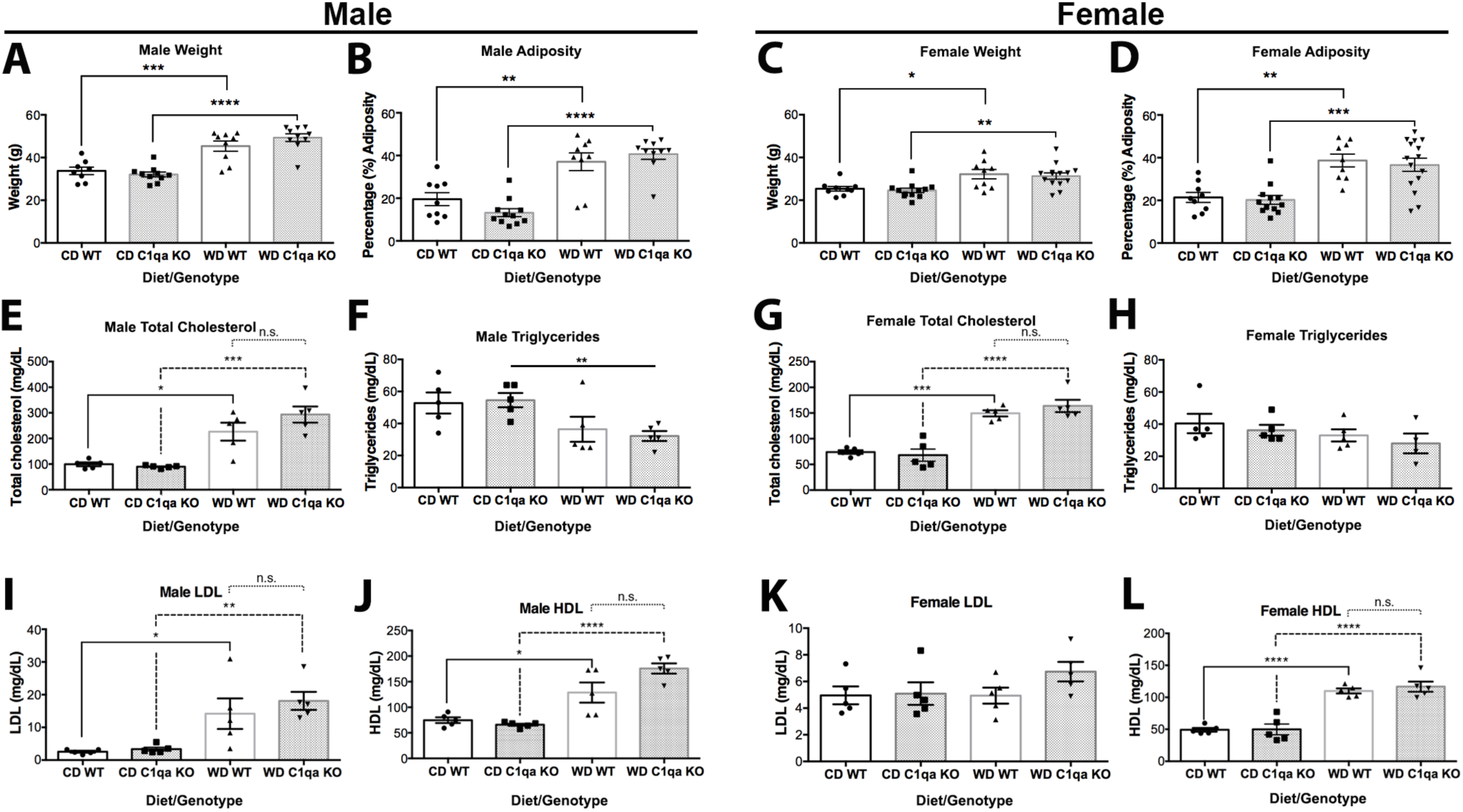
WD-induced obesity and metabolic profiles were not altered by C1QA deficiency. (**A-D**) Weight and adiposity increased in WD-fed mice independent of genotype and sex. (**E-H**) No differences were detected in total cholesterol (E, G) and triglycerides (F, H) between WD-fed WT and *C1qa* KO mice. (**I-L**) LDL and HDL plasma levels in male and female WD-fed mice were not changed by C1QA deficiency. (**M-N**) Fatty acids plasma levels were not altered by C1QA deficiency. Data are presented as mean ± SEM, n ≥ 5, *P < 0.05, **P< 0.01, ***P< 0.001, ****P < 0.0001 by One Way ANOVA with Tukey post hoc test.

**Figure 4.**
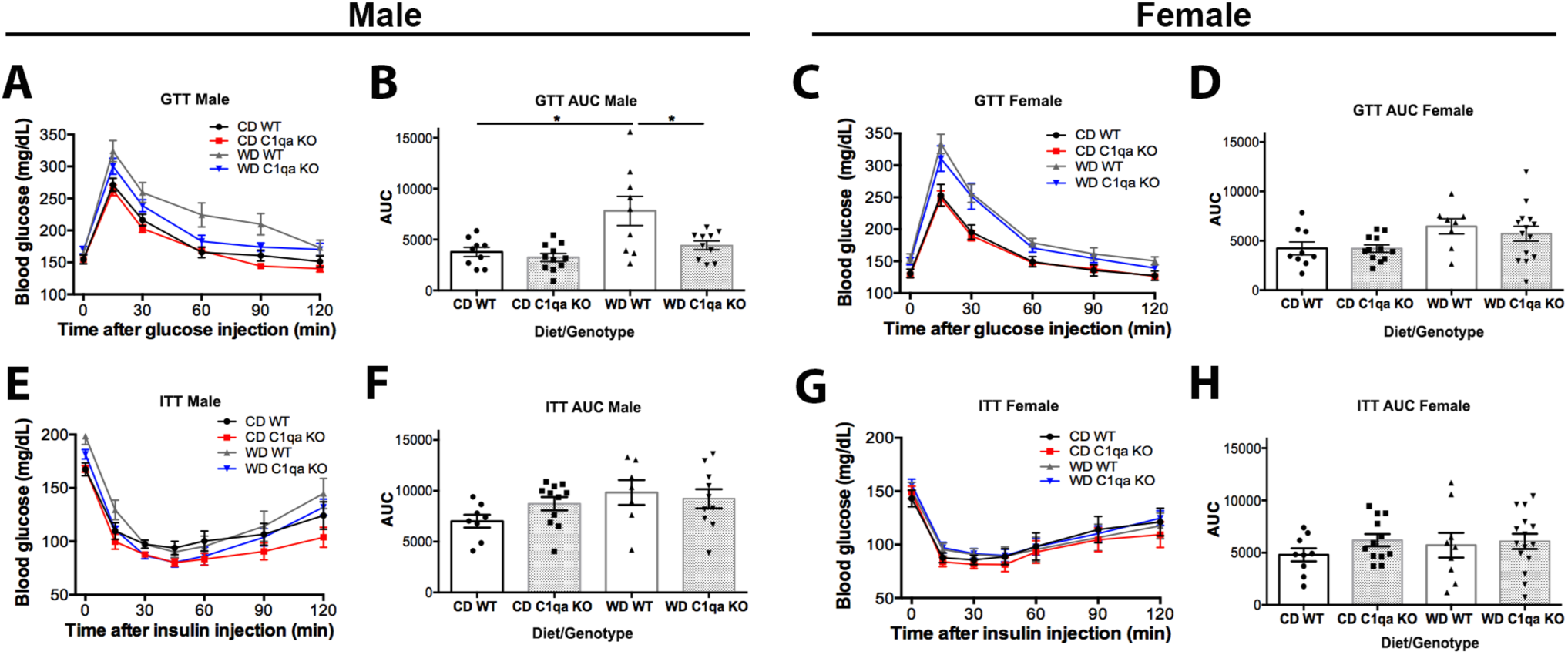
C1QA deficiency did not alter glucose or insulin tolerance in obese mice. (**A-D**) No significant changes were detected in glucose tolerance between genotypes and sex. Only WD-fed male mice showed a moderate impaired glucose tolerance that was prevented by C1QA deficiency (B). (**E-H**) No changes were observed in the insulin tolerance test between groups. Data are presented as mean ± SEM, n ≥ 5, *P < 0.05 by One Way ANOVA with Tukey post hoc test.

Cognitive health was tested in in all mice at 12 mos using a previously validated spontaneous alternation Y-maze task, a measure of working memory (Graham et al., 2019). Male and female CD-fed mice were able to perform the task. In contrast, WD-fed male WT mice showed a significant impairment in their ability to alternate between the arms of the Y-maze indicative of a deficit in working memory (**Fig. 5A-D**). In contrast, male and female *C1qa* KO mice either fed the WD or the CD were able to perform the task (**Fig. 5A-D**). No differences in locomotor activity levels (**Fig. 5E, G**) and grip strength (**Fig. 5F, H**) were found between the experimental groups. These data suggest that although C1QA deficiency did not alter obesity it did prevent deficits in working memory observed in WD-fed male mice.

**Figure 5.**
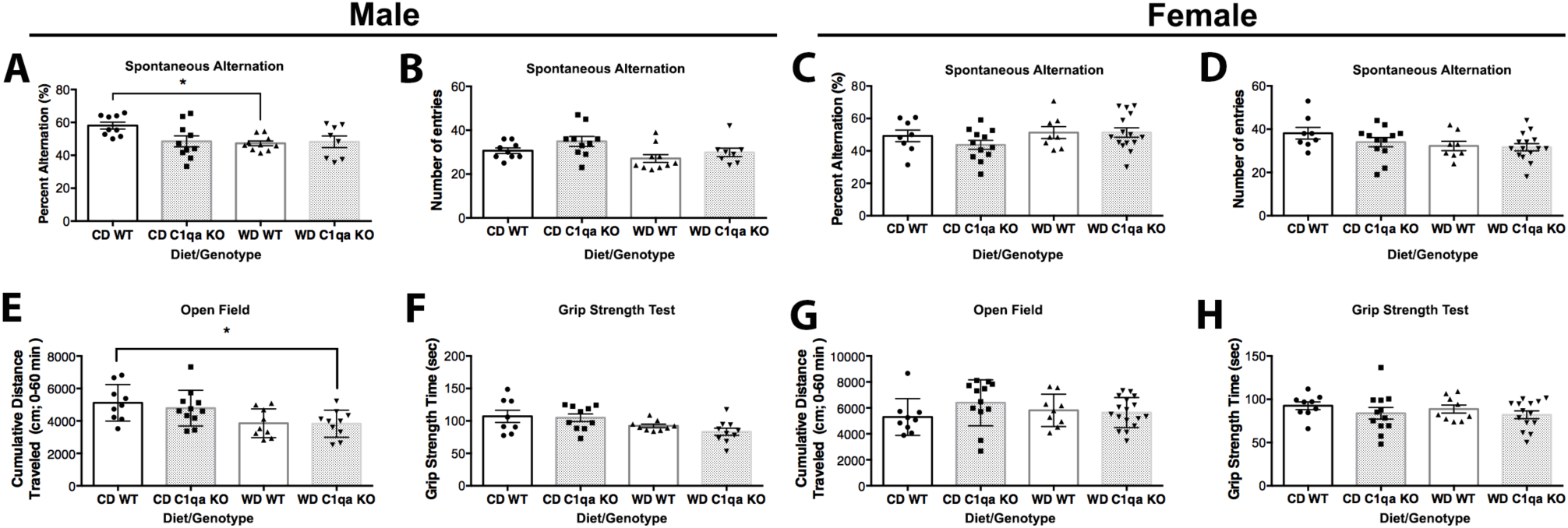
C1QA deficiency prevented WD-induced cognitive deficits in male mice. (**A-B**) WD-fed male mice showed a significant impairment in % correct alternation (A), which was not due to reductions in total activity as measured by total arm entries (B). However, *C1qa* KO WD-fed male mice showed no significant impairment in % correct alternations when compared to CD-fed male mice. (**C-D**) No differences were observed in % correct alternations between CD-fed and WD-fed female mice. (**E, G**) Activity levels in the open field as measured by cumulative distance traveled revealed no differences between groups in both male and female mice. (**F, H**) Grip strength measurements were not different between groups in both male and female mice. Data are presented as mean ± SEM, n ≥ 8, *P < 0.05 by One Way ANOVA with Tukey post hoc test.

The effect of complement activation on cerebrovascular function in aging, diseased brains or obesity is not well studied. Cerebrovascular damage occurs in the aging brain in humans and animal studies (Bell et al., 2010; Montagne et al., 2015; Soto et al., 2015). Importantly, pericyte and basement membrane coverage on vessels is significantly reduced in aged mice and correlates with an increase in *C1qa*-expressing myeloid cells (Soto et al., 2015). Age-dependent cerebrovascular damage is exacerbated by obesity (Tucsek et al., 2014; Graham et al., 2019). As with aging, cerebrovascular damage correlates with increased expression of *C1qa* in myeloid cells (Soto et al., 2015). To determine the potential for complement inhibition to prevent cerebrovascular damage in obesity, we first assessed pericytes number using platelet derived growth factor receptor beta (PDGFRb) immunoreactivity. The number of PDGFRb^+^ cells was significantly reduced in WD-fed WT obese mice compared to CD-fed WT mice (**Fig. 6A-B**). In contrast, there was no significant difference in the numbers of PDGFRb^+^ cells in *C1qa* KO mice fed either the WD or CD. We next assessed laminin, a major component of the basement membrane. While there was a significant decrease in the density of laminin^+^ capillaries in WD-fed WT obese mice when compared to CD-fed mice, no significant difference was found in WD-fed *C1qa* KO obese mice when compared to mice fed the CD (**Fig. 6A, C**). These results suggest C1Q plays a role in obesity-induced cerebrovascular damage.

**Figure 6.**
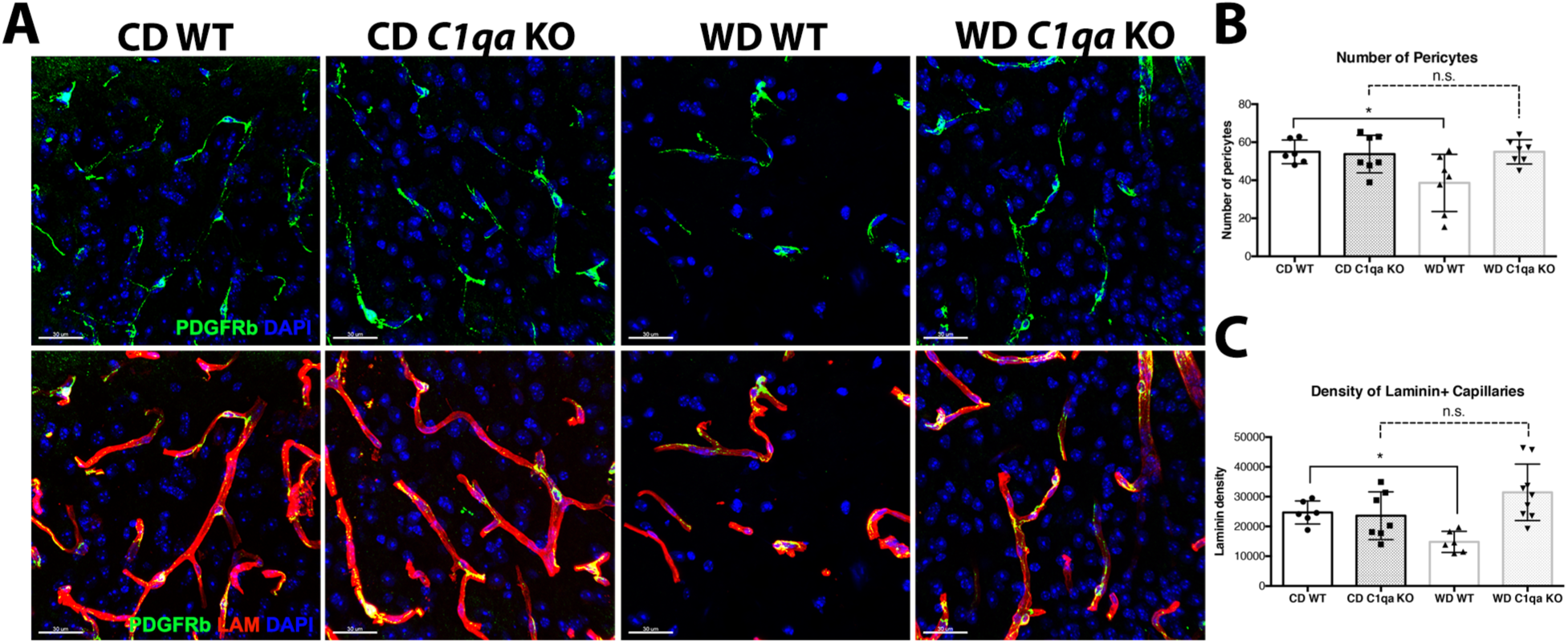
C1QA deficiency prevented WD-diet induced cerebrovascular damage. (**A-C**) WD-induced decrease in PDGFRb^+^ pericytes in WT mice was prevented by C1QA deficiency (A, B). The decrease of LAMININ^+^ capillaries density caused by WD was prevented by C1QA deficiency (A, C). Representative images are from a region of the parietal cortex (above the CA1 region of the hippocampus, see Methods). Data are presented as mean ± SEM, n ≥ 5, *P < 0.05 by One Way ANOVA with Tukey post hoc test. Scale bars: 30μm.

High fat diets, westernized diets and obesity have been shown to induce systemic inflammation, including proinflammatory responses in peripheral tissues and can lead to neuroinflammation and cognitive decline (Ownby, 2010). In this study (**Figs. 1-2**) and our previous studies (Soto et al., 2015; Graham et al., 2016), IBA1^+^ myeloid cells appear to produce the majority of *C1qa* transcript and C1QA protein in the brains of WD-fed obese mice. As reported recently by our group, increased phagocytosis of myelin by microglia is observed in WD-fed obese mice (Graham et al., 2019). Therefore, to determine if C1QA deficiency prevents obesity-induced phagocytic activity of microglia and myelin degradation, brain sections that contained a portion of the CA1 region of the hippocampus, corpus callosum and cortex from 12 mos WD-fed and CD-fed *C1qa* KO and WT mice were immunostained for IBA1 (myeloid cells marker), CD68 (lysosomes/phagosomes) and MBP (myelin basic protein). Images were processed and analyzed using IMARIS (with brain regions batched together, see Methods). There were significantly greater IBA1^+^ surfaces in the WD-fed WT compared to CD-fed WT mice (**Fig. 7A-D, E**). There were also significantly greater CD68^+^ surfaces in WD-fed WT compared to CD-fed WT mice (**Fig. 7A-D, F**), indicative of increased phagocytic activity. Deficiency of C1QA prevented the increase in both IBA1^+^ and CD68^+^ surfaces in WD-fed *C1qa* KO mice (**Fig. 7E, F**).

**Figure 7.**
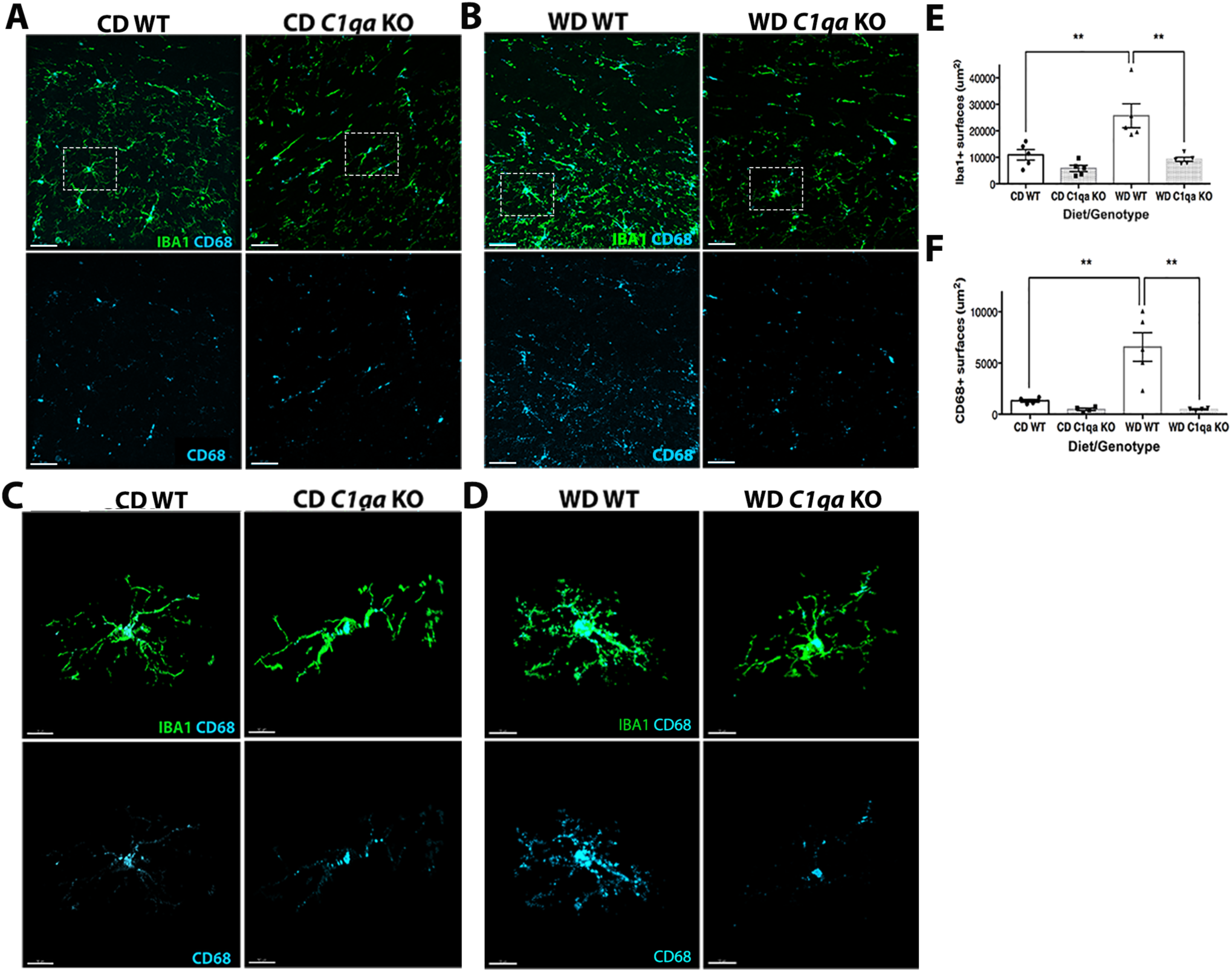
WD-induced myeloid cell activation was prevented by C1QA deficiency. (**A-B**) The increase in the number of IBA1^+^ (A, B, E) and CD68^+^ (A, B, F) surfaces in WD-fed WT was prevented by C1QA deficiency. (**C-D**) Representative examples of IBA1^+^CD68^+^ microglia. Microglia from WD-fed WT mice showed the greatest levels of CD68. Representative images for quantification are shown that contain a portion of the CA1 of the hippocampus, the corpus callosum and the parietal cortex. Data are presented as mean ± SEM, n ≥ 4, **P < 0.01 by One Way ANOVA with Tukey post hoc test. Scale bars: A-B, 30μm; C-D, 10μm.

Phagocytosis of myelin by activated microglia is increased in the presence of complement proteins (DeJong and Smith, 1997). Therefore, myelin positive regions in a portion of the CA1 region of the hippocampus, the corpus callosum and the parietal cortex were assessed using MBP, a major constituent of myelin. Images were again processed and analyzed using IMARIS (see Methods). As expected based on our previous study (Graham et al., 2019), MBP^+^ surfaces in brain sections from WD-fed WT mice were significantly decreased when compared to CD-fed WT mice, particularly in the corpus callosum (**Fig. 8A, B, D**). However, no changes in MBP levels were found in brain sections of WD-fed *C1qa* KO compared to CD-fed WT and *C1qa* KO mice, indicating that deficiency of C1QA prevented the obesity-induced degradation of myelin. Next, the interactions of IBA1^+^ cells with MBP immunostained myelin were assessed. MBP:IBA1 (myelin-myeloid cells) interactions were significantly increased in WD-fed WT mice compared to CD-fed WT mice (**Fig. 8A, B, E**). Colocalization of MBP^+^ surfaces with both CD68^+^ and IBA1^+^ surfaces were commonly observed in WD-fed WT mice but were not observed in WD-fed C1qa KO mice (**Fig. 8C**). These findings suggest C1QA deficiency prevented the phagocytosis of myelin by microglia.

**Figure 8.**
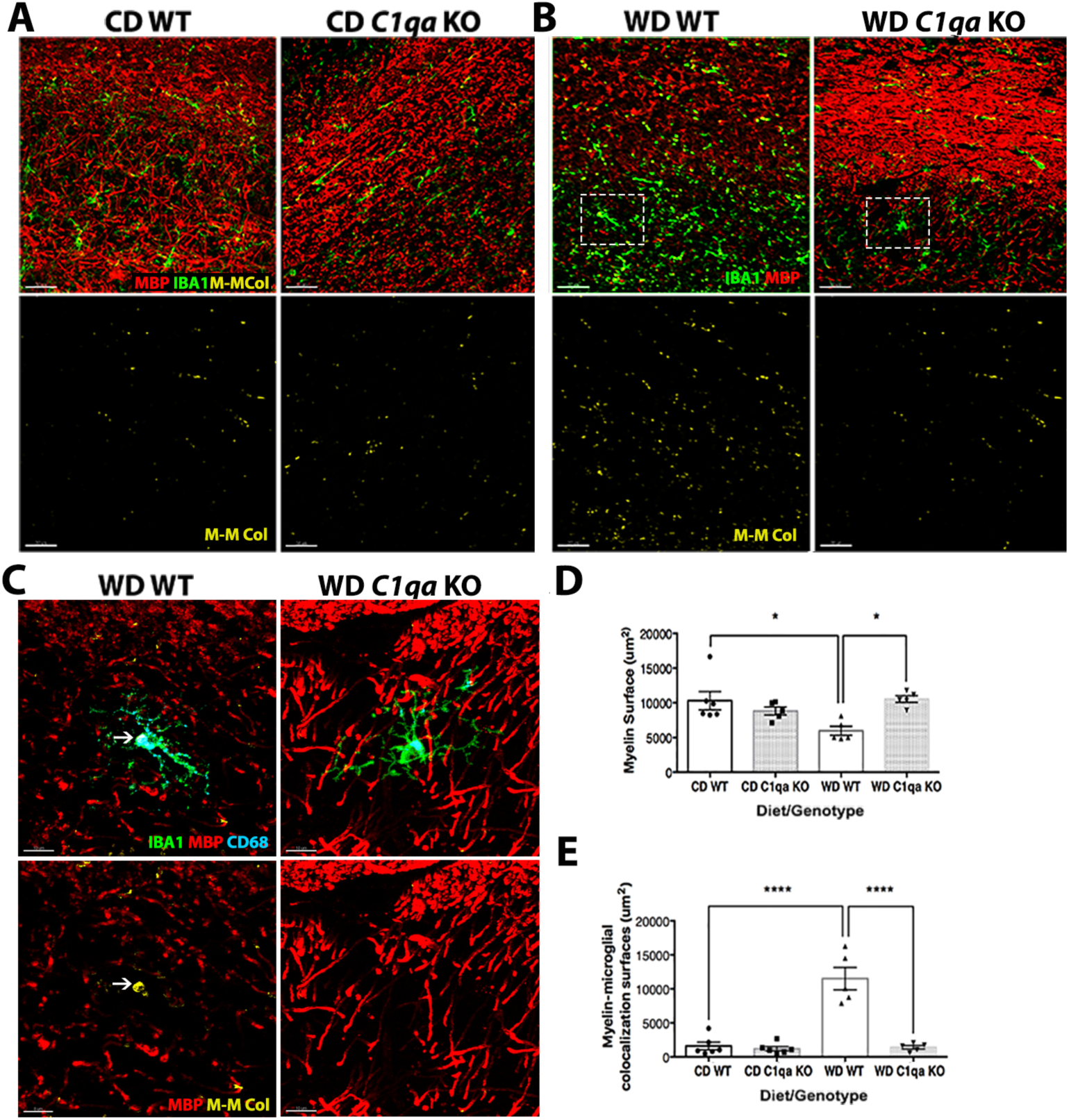
WD-induced myelin loss was prevented by C1QA deficiency. (**A-D**) C1QA deficiency in WD-fed mice prevented the decline of MBP^+^ surfaces. The white arrow (C) indicates an example of CD68^+^ and MBP^+^ surfaces surrounded by IBA1^+^ surfaces, suggesting engulfment. (**E**) C1QA deficiency prevented the increased microglia-myelin interactions observed in WD-fed WT obese mice. Representative images for quantification are shown in (A) and (B) that contain a portion of the CA1 of the hippocampus, the corpus callosum and the parietal cortex. The region of the image within the white boxes in (B) are shown in high resolution in (C). Data are presented as mean ± SEM, n ≥ 4, *P < 0.05, ****P < 0.0001 by One Way ANOVA with Tukey post hoc test. M-M Col = Myelin-Microglia Colocalization. Scale bars: A-B, 30μm; C, 10μm.

## DISCUSSION

The increase in many diet- and obesity-related diseases in western societies, including cardiovascular disease, Type 2 diabetes and dementias have been linked to lifestyle changes. However, little is known about how these changes contribute to long term brain health and increase risk for age-related neurodegenerative disease such as Alzheimer’s disease and related dementias (Elias et al., 2005; Esser et al., 2014). Here, we assessed the effects of globally ablating C1QA, an initiating factor in the classical pathway of the complement cascade, using *C1qa* KO mice. Central components of the classical pathway of the complement cascade, including C1Q and C3, were particularly increased in the corpus callosum of WD-fed compared to CD-fed mice that also presented cerebrovascular dysfunction, microglial activation and white matter degradation. Obesity and high body mass index have been shown to affect other white matter tracks in the brain (Bolzenius et al., 2015; Papageorgiou et al., 2017), but whether complement deposition is present in other white matter tracts in our model is still to be determined.

Activation of the complement cascade occurs in the aging brain and in neurodegenerative diseases (reviewed in (Hammond et al., 2019)). This is the first study to support a damaging role for C1QA in obesity-induced cerebrovascular damage and white matter loss. However, the role of the complement cascade in aging and neurodegenerative diseases has been more widely studied. Inhibition of C1QA prevented synapse loss in aging mice and in mouse models of AD, Parkinson’s disease, and glaucoma (Hammond et al., 2019). However, although inhibition of C3 appeared protective to the aging brain, deficiency of C3 exacerbated some phenotypes in mouse models of AD and glaucoma (Hammond et al., 2019). In our study, to assess the role of C1QA in western diet-induced obesity, we chose to focus on cerebrovascular and white matter structures as well as microglia activity based on findings from two previous studies (Graham et al., 2016; Graham et al., 2019). In the first study, in response to a WD, neuroinflammation was identified as a major consequence of chronic consumption of a WD. A small but significant loss of neurons in the cortex but not the hippocampus was identified (Graham et al., 2016). In the second study, transcriptional profiling supported increased neuroinflammation, as well as cerebrovascular and white matter changes in response to a WD (Graham et al., 2019). In this study, C1QA deficiency prevented pericyte loss and the reduction in laminin density observed in C1QA sufficient WT mice fed the WD. Additional experiments are still required to determine the degree to which C1QA deficiency impacts blood brain barrier integrity and function. The effect of complement deficiency on neuronal numbers as well as neuronal structures such as synapses and axons in obese brains is also still to be determined. However, this study further supports the complex and important role of the complement cascade in aging, obesity and neurodegenerative diseases.

The pathologies described here as a result of WD-induced obesity, that were prevented by C1QA deficiency, are similar to those observed in cerebrovascular-relevant dementias – often grouped as vascular contributions to cognitive impairment and dementia (VCID). The pathologies include small vessel disease, neurovascular dysfunction and white matter hyperintensities or diffuse demyelination (de Leeuw et al., 2001; van Norden et al., 2011; Alber et al., 2019). The effect of complement activation on cerebrovascular health in aging, diseased brains or obesity is not well studied. Cerebrovascular dysfunction occurs in the aging brain in humans and animal studies (Bell et al., 2010; Montagne et al., 2015; Soto et al., 2015). Importantly, pericyte and basement membrane coverage on vessels is significantly reduced in aged mice and correlates with an increase in *C1qa*-expressing myeloid cells (Soto et al., 2015). Age-dependent cerebrovascular decline is exacerbated by obesity (Tucsek et al., 2014; Graham et al., 2019). As with aging, cerebrovascular decline correlates with increased expression of *C1qa* in myeloid cells (Soto et al., 2015). C3 deposition has been reported on cerebral vessels of AD patients, suggesting a possible role of complement in cerebrovascular decline (Shi et al., 2019). Furthermore, complement synthesis is increased in activated microglia preceding blood brain barrier dysfunction in an inducible rat model of neurotoxicity, suggesting complement proteins can directly damage the cerebrovasculature (Lynch et al., 2004). In fact, it is thought that the cerebrovasculature is particularly susceptible to complement components as it can be exposed to these proteins that are produced in the brain and/or circulating in the blood. *In vitro*, complement receptors such as the C3aRs and C5aRs are expressed in small numbers on endothelial cells and can induce signaling cascades that are distinct from leukocytes (Schraufstatter et al., 2002; Hernandez et al., 2017). C1Q can bind endothelial cells directly causing increased expression of adhesions molecules such as intracellular adhesion molecule-1 (ICAM-1) and vascular cell adhesion molecule-1 (VCAM-1) (Storini et al., 2005). Therefore, our data supports targeting members of the complement cascade as a viable strategy to alleviate symptoms in chronic brain disorders that include a significant cerebrovascular component.

White matter changes that appear to have occurred as a result of damaging actions by microglia were prevented by C1QA deficiency in obese mice. However, the initiating factors of these damaging responses are not clear. Previous studies have shown that microglia play an important role in the degradation of myelin through myelin turnover. Phagocytosis of myelin by activated microglia is increased in the presence of complement proteins (DeJong and Smith, 1997). Interestingly, one study showed that specifically targeting pericytes was sufficient to induce white matter dysfunction (Montagne et al., 2018), indicating that pericyte ‘stress’ in response to chronic consumption of a WD may be an early and important event in our obese mice. Neurovascular unit breakdown and cerebral small vessel disease are known to increase damaging neuroinflammatory responses by microglia and astrocytes (del Zoppo, 2009; Soto et al., 2015). Astrocyte reactivity could directly impact myelin turnover as it can cause a breakdown in astrocyte-oligodendrocyte gap junctions leading to a failure of remyelination (Sharma et al., 2010; Markoullis et al., 2014). In addition, astrocyte reactivity may cause a breakdown in astrocyte-pericyte or astrocyte-endothelial cell interactions leading to cerebrovascular damage (Abbott, 2002; Zhao et al., 2015). Given the suggested roles in neuronal dysfunction, cerebrovascular breakdown and white matter damage, targeting the complement cascade remains an important area to study for brain disorders that present multiple endophenotypes, as is the case for many dementias.

Global deletion of the *C1qa* gene prevented cognitive decline and the pathological changes to the cerebrovasculature and white matter found in the brain of obese mice. However, it is not clear whether the protection is due to the lack of C1QA in the brain or systemically. Assessment of phenotypes relevant to metabolic syndrome such as weight, adiposity and blood cholesterol were not significantly affected by C1Q deficiency (**Figs. 3 and 4**). However, a detailed examination of lipids and metabolites in the blood or the brain (e.g. by lipidomics and metabolomics) was not performed meaning we cannot rule out that the protection to brain structures may be driven by changes to lipid or metabolite species in C1QA deficient compared C1QA sufficient mice fed the WD. The C1QA-mediated protection to brain structures may also be working at the level of changes to systemic inflammation. High fat diets, westernized diets and obesity have been shown to induce systemic inflammation, including proinflammatory responses in peripheral tissues and can lead to neuroinflammation and cognitive decline (Ownby, 2010). Systemic inflammation has also been shown to damage the microvasculature of the brain in aging and dementia (Grammas and Ovase, 2001; Grammas et al., 2006). However, in this study, obesity-induced cerebrovascular damage and white matter loss was prevented by C1QA deficiency without altering the weight and metabolic profiles of these mice which were similar to the WD-fed WT mice. These findings suggest the protection mediated by C1QA deficiency may be due to inhibition of complement-expressing microglial cells directly. Although circulating complement components account for approximately 4% of blood proteins, in young, aging diseased brains, complement components are also synthesized by multiple cell types including myeloid cells, astrocytes, neurons and endothelial cells (Hammond et al., 2019). In the aging brain, microglia and astrocytes are thought to be a major source of complement proteins (Fonseca et al., 2017). Our data suggest microglia are the major producers of C1QA (**Figs. 1, 2**) in obese brains. Interestingly, previous studies in our laboratory support infiltration of peripherally-derived myeloid cells (monocytes/macrophages and neutrophils) into the brain during aging that is increased in diet-induced obese mice (Yang, 2019) indicating both resident and peripherally-derived cell types might contribute to increased complement proteins in brains of obese mice. Therefore, although we refer to the IBA1^+^ and C1QA^+^ cells in the brain as microglia, we cannot rule out a contribution from peripherally-derived IBA1^+^ myeloid cells. Conditionally ablating components of the complement cascade such as C1Q and C3 in specific cell types will be needed to precisely determine the source of complement. However, although more work is required, this study provides further support for exploring complement inhibition as viable route to preserve brain health throughout aging.

## Authors’ Contributions

LCG and GRH conceived the project. LCG, HEK and IS performed all of the experiments and data analysis. LCG, HEK, IS and GRH wrote the manuscript and all authors reviewed and approved the final version.

## Acknowledgements

The authors thank: Drs. Simon John, Mimi DeVries and Jeffrey Harder for western diet development, Keating Pepper for help with IMARIS, Dr. Stacey Rizzo and Laura Anderson for behavioral phenotyping, Harriet Williams for help with creation and validation the *C1qa^-/-^* mice, and John West and Yu-Chien Wu for help with the imaging analysis. This work was funded by RF1 AG051496 (GRH).

## Supplemental figures and legends

**Figure S1.**
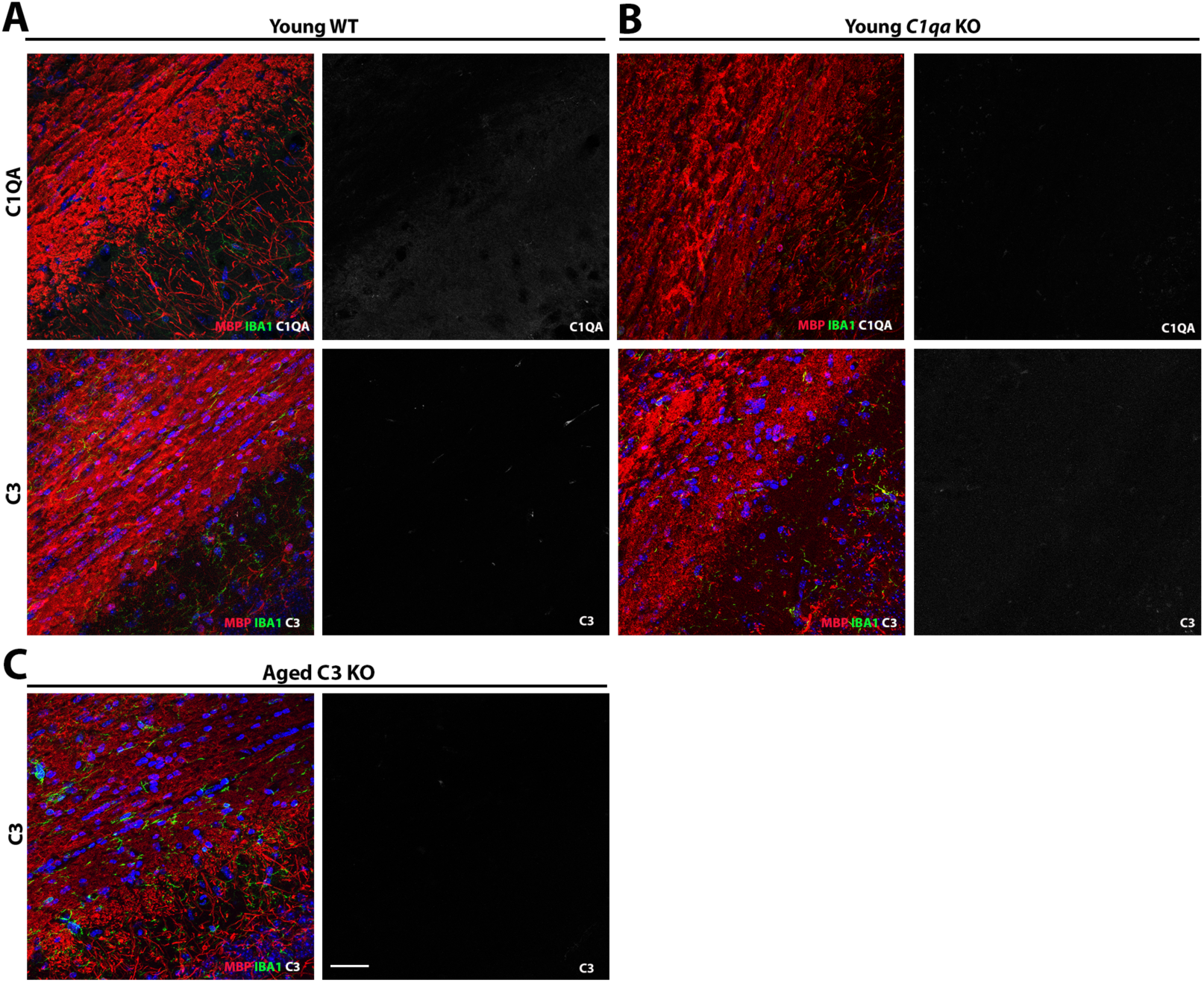
C1QA and C3 immunoreactivity in young WT and *C1qa* or *C3* knockout mice. (**A**) C1QA (upper panel) and C3 (lower panel) immunoreactivity in the young WT corpus callosum is low or absence. (**B**) No C1QA immunoreactivity is found in the *C1qa KO* mice. (**C**) No C3 immunoreactivity is found in the *C3 KO* mice. Scale bars: 40μm.

**Figure S2.**
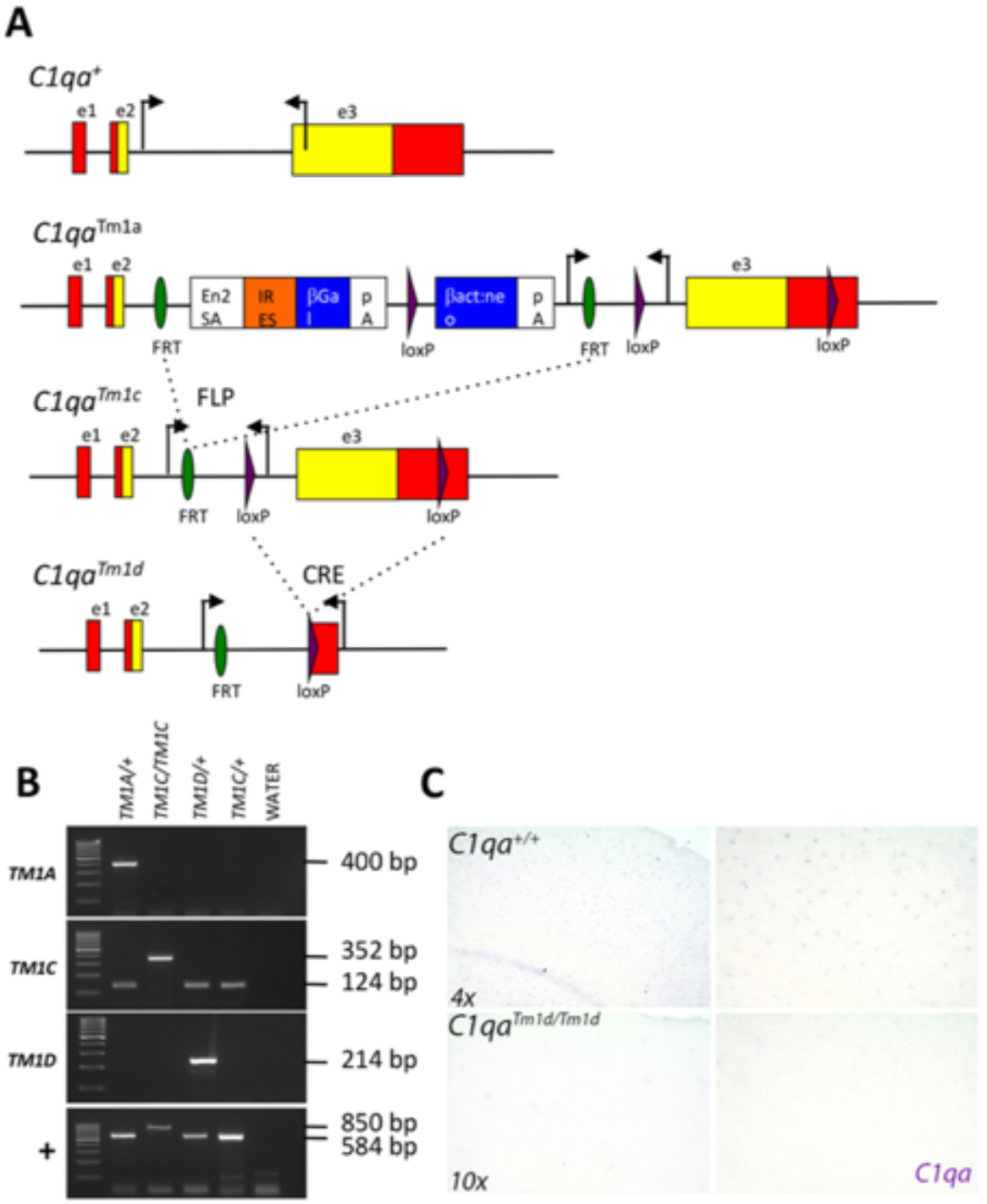
Creation and validation of the *C1qa* KO mouse. A conditional *C1qa* mouse was created as part of the Knockout Mouse Project (KOMP) at The Jackson Labs (https://www.jax.org/research-and-faculty/tools/knockout-mouse-project). (**A**) Representation of the *C1qa* alleles that are available including *C1qa^Tm1a^* (LacZ reporter), *C1qa^Tm1c^* (floxed allele), *C1qa^Tm1d^* (null allele). As has been shown for other KOMP alleles, the *Tm1a* allele is not capable of reporting *C1qa* expression using the β-galactosidase assay. Mating the *C1qa^Tm1a^* mice to mice carrying FLP recombinase creates the *C1qa^Tm1c^* floxed allele. Mating the *C1qa^Tm1c^* mice to mice carrying CRE recombinase creates the *C1qa^Tm1d^* null allele. Cell-specific ablation of *C1qa* can be achieved using a cell-specific Cre line. The locations of genotyping primers are shown as inward facing arrows. (**B**) Examples of genotyping assays for the different alleles and band sizes. Primers pairs in this example were as follows. *C1qa^+^*: F – CCGGAAGAAAAGACATCCTG; R – CTTTCACGCCCTTCAGTCCT. C1qa*^Tm1a^*: F – GTGGTTTGTCCAAACTCATCAA; R – TCTCTGAGCCTCTGCTTCAA. *C1qa^Tm1c^*: F – GGACGAGAGGGGAGGAGTTA; R – TTAGGACCCTTTGGCACAAC. *C1qa^Tm1d^*: F – CCGGAACCGAAGTTCCTATT; R – AGACGGGGATCGTTTATTCC. Note that the *C1qa^+^* primers amplify a larger product in mice carrying the *C1qa^Tm1c^* allele. Standard PCR conditions were used with an annealing temperature of 59.3°C for *C1qa^+^*, *C1qa^Tm1b^* and *C1qa^Tm1d^* and 61.0°C for *C1qa^Tm1c^*. (**C**) RNA in situ hybridization to visualize *C1qa* transcripts in brains sections from *C1qa^+/+^* and *C1qa^Tm 1d/Tm1d^* mice. As expected, *C1qa* expression is absent in *C1qa^Tm1d/Tm1d^* (*C1qa* KO) mice.

